# MuLan-Methyl - Multiple Transformer-based Language Models for Accurate DNA Methylation Prediction

**DOI:** 10.1101/2023.01.04.522704

**Authors:** Wenhuan Zeng, Anupam Gautam, Daniel H. Huson

## Abstract

Transformer-based language models are successfully used to address massive text-related tasks. DNA methylation is an important epigenetic mechanism and its analysis provides valuable insights into gene regulation and biomarker identification. Several deep learning-based methods have been proposed to identify DNA methylation and each seeks to strike a balance between computational effort and accuracy. Here, we introduce MuLan-Methyl, a deep-learning framework for predicting DNA methylation sites, which is based on five popular transformer-based language models. The framework identifies methylation sites for three different types of DNA methylation, namely N6-adenine, N4-cytosine, and 5-hydroxymethylcytosine. Each of the employed language models is adapted to the task using the “pre-train and fine-tune” paradigm. Pre-training is performed on a custom corpus of DNA fragments and taxonomy lineages using self-supervised learning. Fine-tuning aims at predicting the DNA-methylation status of each type. The five models are used to collectively predict the DNA methylation status. We report excellent performance of MuLan-Methyl on a benchmark dataset. Moreover, we argue that the model captures characteristic differences between different species that are relevant for methylation. This work demonstrates that language models can be successfully adapted to applications in biological sequence analysis and that joint utilization of different language models improves model performance. Mulan-Methyl is open source and we provide a web server that implements the approach.

**Key points:**

- MuLan-Methyl aims at identifying three types of DNA-methylation sites.
- It uses an ensemble of five transformer-based language models, which were pre-trained and fine-tuned on a custom corpus.
- The self-attention mechanism of transformers give rise to importance scores, which can be used to extract motifs.
- The method performs favorably in comparison to existing methods.
- The implementation can be applied to chromosomal sequences to predict methylation sites.

## Introduction

DNA methylation is an important biological process. It facilitates epigenetic regulation of gene expression, is associated with various medical disorders [1, 2, 3], and has other applications, such as a marker in metagenomic binning [4]. While DNA methylation is a dynamic process, existing machine-learning techniques are able to predict DNA methylation states from genomic sequence with some degree of accuracy.

There are several types of DNA methylation that differ by which methyl group is attached to which type of nucleotide in the sequence. Here, we focus on 6-methyadenine (6mA), 5-hydroxymethylcytosine (5hmC), and 4-methylcytosine (4mC) methylation [5, 6, 7]. Different organisms exhibit different patterns of methylation and this gives rise to the computational problem of predicting the location of methylation sites for a given genome sequence. While much algorithmic work has been done on the question, recent work has focused on the application of deep learning methods [8, 9]. However, there is room for improvement of accuracy and comprehensiveness.

A large number of papers address the problem of identifying methylation sites, however, most of them focus on specific form of modification [10, 11, 12, 13, 14, 15, 16, 17, 18, 19, 20, 21, all three types of methylation mentioned above [30, 31, 32, 33, 34], in particular iDNA-MS, iDNA-ABT, and iDNA-ABF. The database presented in [31**?**] is now widely used as a benchmark dataset for assessing model performance [21, 23, 32, 33, 34].

While different deep-learning based methods all address the same goal, they differ in the details of the features employed and the model structure. Input features include an encoding of the sequence, of course, but may also include biochemical properties [10, 12], or a DNA molecular graph representation [22], say. Utilized model structures include Convolutional Neural Networks (CNN), Graph Convolutional Neural Networks (GCN), Bidirectional Encoder Representation from Transformers (BERT) [35], and other types of machine learning algorithms. The specific choices made during feature engineering and model selection determine performance and are key to proposing a new framework.

Here, we phrase DNA methylation-site detection as a Natural Language Processing (NLP) problem and propose a novel framework to address it. Previous studies for identifying methylation sites usually use BERT, a classic NLP approach, or, in the context of DNA sequences, the variant DNABERT [36], either as a model that accepts embeddings from Word2vec, or as an encoder that generates embeddings for input to a deep neural network [37, 23, 32, 25, 33]. Only few published approaches aim at predicting multiple DNA modification sites. Moreover, many do not use taxonomic information as explicit features, although the taxonomic identity of an organism has an impact on DNA methylation [38]. Here we address both shortcomings by providing a new framework that uses a set of collectively training language models, including, but not limited to BERT, to predict three types of methylation sites from DNA sequences and taxonomic information.

Combining the transformer-based language model BERT with the “pre-train and fine-tune” paradigm has become the method of choice in NLP applications. In the pre-training step, self-supervised learning of the Masked Language Modelling (MLM) task and the Next Sequence Prediction (NSP) task are usually performed on a corpus consisting of Wikipedia and books. This allows the transformer-based language model to capture the semantics of text input and contextual information exceptionally well. Transformer-based language models dynamically learn the input’s representation through a multi-head self-attention mechanism [39] and this leads to an improvement of prediction over classification models constructed using static embedding approaches [40]. The fine-tuning step involves supervised training of the pre-trained language model to adapt to specific downstream tasks, here the prediction of three different types of methylation sites. Using BERT as a starting point, and then varying the network architecture and parameters, one can obtain five different language models, [41, 42, 43, 44, 45]. By pre-training on a domain-specific custom corpus, BERT can be adapted to a specific application scenario [46, 47, 48, 49]. While the analysis of DNA sequences can be considered an application of NLP, using language models that are trained on human languages will not do well at capturing nucleotide rules. To address this, several approaches, such as BERTax, DNABERT and LOGO [50, 36, 51], use large amounts of genomic sequence, instead of Wikipedia, as a corpus or similar structure.

The main aim of this paper is to introduce MuLan-Methyl, a novel deep-learning framework that combines five transformer-based language models to collectively predict sites for three different types of methylation (see Figure 1A). In this approach, each methylation-site sample is written as a sentence that represents the surrounding DNA sequence and the taxonomic identity of the corresponding genome. The output of our model is based on the average of the prediction probabilities obtained by five transformer-based language models, namely BERT [35], DistilBERT [41], ALBERT [45], XLNet [43] and ELECTRA [44].

**Fig. 1.**
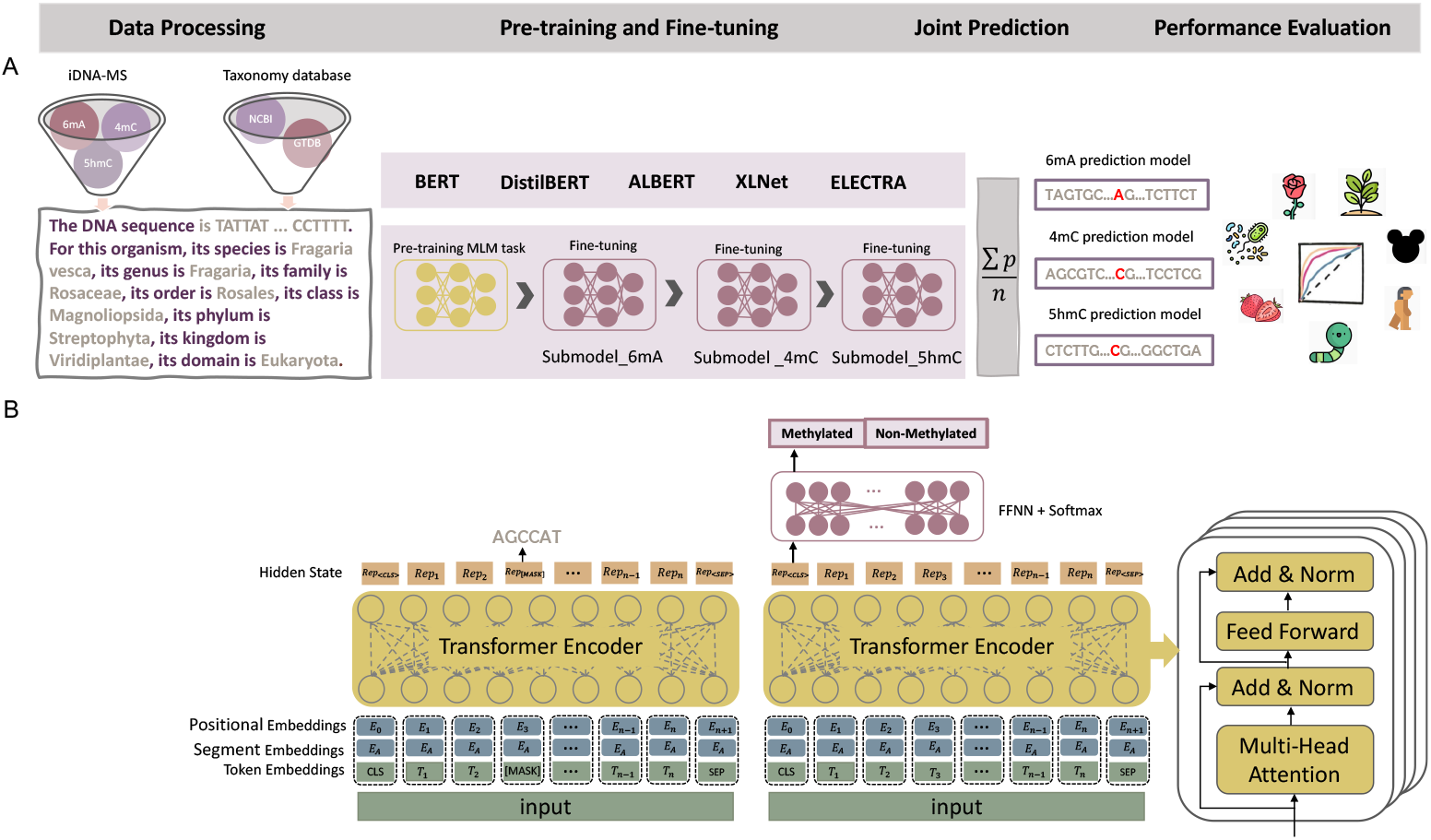
The MuLan-Methyl workflow. (A) The framework employs five fine-tuned language models for joint identification of DNA methylation sites. Methylation datasets (obtained from iDNA-MS) were processed as sentences that describe the DNA sequence as well as the taxonomy lineage, giving rise to the processed training dataset and the processed independent set. For each transformer-based language model, a custom tokenizer was trained based on a corpus of the processed training dataset and taxonomy lineage data. Pre-training and fine-tuning were both conducted on each methylation-site specific training subset separately. During model testing, the prediction of a sample in the processed independent test set is defined as the average prediction probability of the five fine-tuned models. We thus obtained three methylation type-wise prediction models. We evaluated the model performance on the genome type that contained in the corresponding methylation type-wise dataset, respectively. In total, we evaluated 17 combinations of methylation types and taxonomic lineages. (B) The general transformer-based language model architecture for pre-training and fine-tuning. The model was pre-trained using the masked language modeling (MLM) task and then fine-tuned on the methylation type-wise processed training datase2t.

Each of the five language models is trained according to the “pre-train and fine-tune” paradigm. For this, we used a custom corpus that contains the processed training dataset and taxonomic lineage information downloaded from NCBI [52] and GTDB [53]. For each language model, we trained a custom tokenizer on the custom corpus, using the same configuration as the model’s default tokenizer. We use a customized tokenizer to ensure that the represented DNA sequences and taxonomic information associated with each sample is captured effectively. Each language model was pre-trained by training the MLM task on the processed training dataset. We then obtained the 6mA model by fine-tuning the pre-trained language model using the 6mA training dataset. Next, the 4mC prediction model was obtained by fine-tuning the 6mA prediction model using the 4mC training dataset. Finally, the 5hmC prediction model was obtained by fine-tuning the 4mC prediction model using the 5hmC training dataset. In addition, we compared the performance of all models contained in MuLan-Methyl.

A main contribution of this work is that we use both DNA sequence and taxonomic identity as explicit features in the model. Using the iDNA-MS [31] independent test set as a benchmark, our approach shows improved performance over previous methods, especially for certain genomes. MuLan-Methyl is capable of making accurate predictions for genomes whose taxonomy lineage is not present in the training dataset. The interpretability of MuLan-Methyl facilitates the discovery of DNA motifs that are associated with DNA methylation and potential correlations between specific methylation sites and taxonomic lineages.

To the best of our knowledge, this is the first application in biology that achieves improved prediction performance by integrating multiple transformer-based language models. This shows that adding features to a model is not the only way to improve the accuracy of predictions.

## MATERIALS AND METHODS

### Data processing

#### Data collection

We downloaded a DNA methylation dataset from [54]. This is an open resource that was published with the iDNA-MS method [31] and is widely used for benchmarking. The dataset contains three main types of DNA methylation sites, namely 6mA, 4mC and 5hmC, across 12 genomes (one bacteria and 11 eukaryotes), in total 250,599 positive samples. In addition, the dataset provides the same number of non-methylation sequences as negative samples.

The dataset is partitioned into a training set and an independent test set at a 1:1 ratio. The training dataset provides samples for methylation type 6mA present in 11 different species. In more detail, the numbers are 53,800 for *T. thermophile*, 15,937 for *A. thaliana*, 9,168 for *H. sapiens*, 8,608 for *Xoc. BLS256*, 5,596 for *D. melanogaster*, 3,981 for *C. elegans*, 3,033 for *C. equisetifolia*, 1,893 for *S. cerevisiae*, 1,690 for *Tolypocladium*, 1,551 for *F. vesca* and 300 for *R. chinensis*. The 4mC methylation type is present in four species, where the numbers of samples are 7,899 for *F. vesca*, 7,664 for *Tolypocladium*, 990 for *S. cerevisiae*, and 183 for *C. equisetifolia*. Finally, the numbers of samples for the type 5hmC are 1,840 for *M. musculus* sequences and 1,172 for *H. sapiens*. More detailed statistics are provided in Supplementary Table S1.

Each sample is a DNA segment of length 41, which is centered on an experimentally-verified methylation site, in the case of a positive sample.

#### Dataset preparation

We processed each sample (DNA sequence of length 41) as follows. Using a sliding window of length 6, we extracted 36 = 41 *−* 6 + 1 individual 6-mers from the DNA sequence, and embed these into a sentence, together with a description of the taxonomic lineage of the corresponding organism, which was phrased as follows: “For this organism, its species is *species*, its genus is *genus*, its family is *family*, its order is *order*, its class is *class*, its phylum is *phylum*, its kingdom is *kingdom*, its domain is *domain*.” We refer to a set of sentences obtained from a set of samples as a “processed dataset.” The full processed training dataset, containing all three types of methylation sites, was put into the custom corpus. For purposes of fine-tuning, both the processed training dataset and the processed independent test set were split into three sets by methylation type.

#### Corpus generation

We require a custom corpus for pre-training each language model to allow the model to learn and capture domain-specific words, which are not contained in a text corpus such as Wikipedia. The custom corpus contains the processed training dataset, as mentioned above. In addition, to enable the language to detect words about taxonomy, we incorporated all taxonomic lineages from the NCBI and GTDB taxonomies that are not already contained in the training datasets. In total, the corpus contains 2,440,894 sentences and uses a vocabulary of 25,000 words.

#### External dataset

We downloaded DNA methylation data published with the Hyb4mC method [16] and with the i6mA-pred method [55], respectively. As this “external” data is not contained in the our training or independent datasets, nor do the associated taxonomic lineages coincide, it is ideal for evaluating the performance of MuLan-Methyl more broadly. In more detail, this data consists of samples (DNA sequences of length 41) representing 320 4mC-site sequences in *E. coli*, 1,926 4mC-site sequences in *G. pickeringii*, and 880 6mA-site sequences in *Oryza sativa* L., each with the same number of negative samples, respectively.

### Training transformer-based language models

We pre-trained and fine-tuned five transformer-based language models. In the following, we first describe the architecture of each of the five employed language models. We then discuss the details of the training process for the first method, BERT, including tokenization, pre-training, and fine-tuning (see Figure 1B). The other four languages are trained in a similar way.

All code is written in Python 3.10, using the Pytorch and the Huggingface Transformers library [56]. The experiments were run on a Linux Virtual Machine (Ubuntu 20.04 LTS) equipped with 4 GPUs provided by de.NBI (flavor: de.NBI RTX6000 4 GPU medium).

#### Transformer-based language models

Our approach uses five transformer-based language models, which we introduce in the following.

1. BERT is capable of modelling bidirectional contexts, using denoising and autoencoding-based pre-training. The transformer architecture of BERT_base_ uses 12 layers in the encoder stack, 768 hidden units for feedforward networks and 12 attention heads; it has 110M parameters in total.
2. A distilled version of BERT, DistilBERT, is obtained by decreasing the number of layers. It has 40% the size of BERT and is 60% faster, while only being 3% less accurate.
3. ALBERT adopts a cross-layer parameter sharing technique for 12 transformer encoder blocks and imports embedding factorization between vocabulary and the hidden layer in order to reduce the parameter size of BERT.
4. XLNet uses an innovative pre-training step; its generalized autoregressive pre-training method enables learning bidirectional contexts by maximizing the expected likelihood over all permutations of the factorization order, overcoming the issues caused by BERT’s neglect of dependency between masked positions.
5. Using a different architecture, ELECTRA trains two transformer models; a generator replaces tokens in a sequence and a discriminator tries to identify which tokens were replaced by the generator, instead of training on the MLM task.

#### Custom tokenizer

A tokenizer must be used to convert samples into the format that is expected by the transformer block of a language. In our study, such a tokenizer is obtained by training the language’s default tokenizer on our custom corpus. Once trained, the tokenizer can capture any sample represented by a sentence consisting of 6-mer DNA words and a textual description of taxonomic lineage.

After tokenization, each input sample is represented by a list of tokens, starting and ending with special tokens [CLS] and [SEP], respectively, and padded to a length of 100 using padding tokens [PAD].

#### Model pre-training

The BERT language model is pre-trained by performing unsupervised training of the MLM task on the custom corpus. Pre-training was conducted on the model using an architecture that is the same as *bert-base-uncase*, but with setting the embedding size of input to 25,000 to match the vocabulary size of the corpus.

During training of the MLM task, 15% of all WordPiece tokens of a sample are selected at random as masking candidates. Of these, 80% are replaced by a special token [MASK] and 10% are replaced by a random token. Then the original tokens are predicted.

Pre-training was conducted using 8 epochs, a batch size of 64 per GPU, and a learning rate of 5e-4, which is achieved after 100 steps of warmup.

#### Model fine-tuning

Fine-tuning is performed for each of the three methylation-site types separately, and so the processed training dataset is split into three training subsets, 6mA, 4mC and 5hmC, listed in order of decreasing size. Each training subset is split into a training set and a validation set at a ratio of 8:2. The target model used to be fine-tuned depended on the subset’s size. First, for the 6mA subset, we simply fine-tuned the pre-trained language model that was trained on the custom corpus. Second, the 4mC fine-tuned model was then obtained by fine-tuning the 6mA fine-tuned model. Finally, 5hmC fine-tuned model was obtained by fine-tuning the 4mC fine-tuned model. We fine-tune the fine-tuned models in this way to make the predictions more accurate on the smaller training subsets.

In all three cases, fine-tuning was performed using an early-stopping strategy, with a maximum of 32 epochs, a batch size of 64 per GPU, and a learning rate of 1e-5, which is achieved after 100 steps of warmup.

### Multi-language model

For each of the three types of methylation sites, five language models are trained and then the MuLan-Methyl framework integrates these, computing prediction probabilities that are obtained by averaging over the probabilities returned by the five models.

### Interpretability of MuLan-Methyl

Transformer-based language models learn different and distant dependencies in the input, by virtue of the multi-head self-attention mechanisms that are present in each encoding layer. For example, BERT contains 12 encoder layers containing 12 attention heads each. For one layer, the multi-head self-attention can be described as

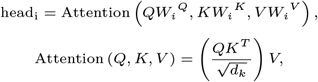

Here, the *i*_*th*_ single attention head is computed as

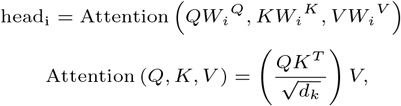

where the projections represent parameter matrices 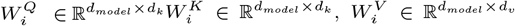, and 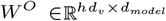. *Attention* = *{a*_*ij*_ *}* is a scoring matrix, in which *a*_*ij*_ denotes the attention weight that the *Query* token *t*_*i*_ gets from then *Key* token *t*_*j*_. This matrix is widely used for representing and exploring the binding between tokens [49, 33, 57].

While the language models are fine-tuned on the methylation-sites prediction task, in the last layer of our model, a softmax function that acts as a classifier is placed on the special token [CLS] that is present at the beginning of each input sentence.

For each token, we sum the attention weights assigned to [CLS] over the 12 heads and regard this as the token’s contribution to sample prediction.

To analyze the impact of the DNA sequence of a sample on the taxonomic lineage of the sample, we extract the attention weights assigned by the DNA tokens to the taxonomic hierarchy tokens.

Note that the WordPiece algorithm, which is used by the tokenizer employed in BERT, DistilBERT and ELECTRA, provides word-wise tokens, so it makes sense to view the attention weights of tokens as contribution scores.

Here we conduct the above computation on the three fine-tuned models of each methylation type in MuLan-Methyl, respectively. The token importance score for MuLan-Methyl is obtained as the average score achieved on each of the three site-specific models.

## RESULTS

### Comparison with encoders from language models

To illustrate the effectiveness of the approaches we proposed for training language models for DNA-based applications, we compared the encoder of our pre-trained language model with that of both BERT and DNABERT (see Figure 2A). Each pre-trained language model was applied to 10% of the positive DNA sequences in the independent test set, obtaining their sentence representation by extracting the embedding of [CLS], with a dimension of (1, 768). The samples were then clustered and visualized using UMAP, colored by taxonomic lineage.

**Fig. 2.**
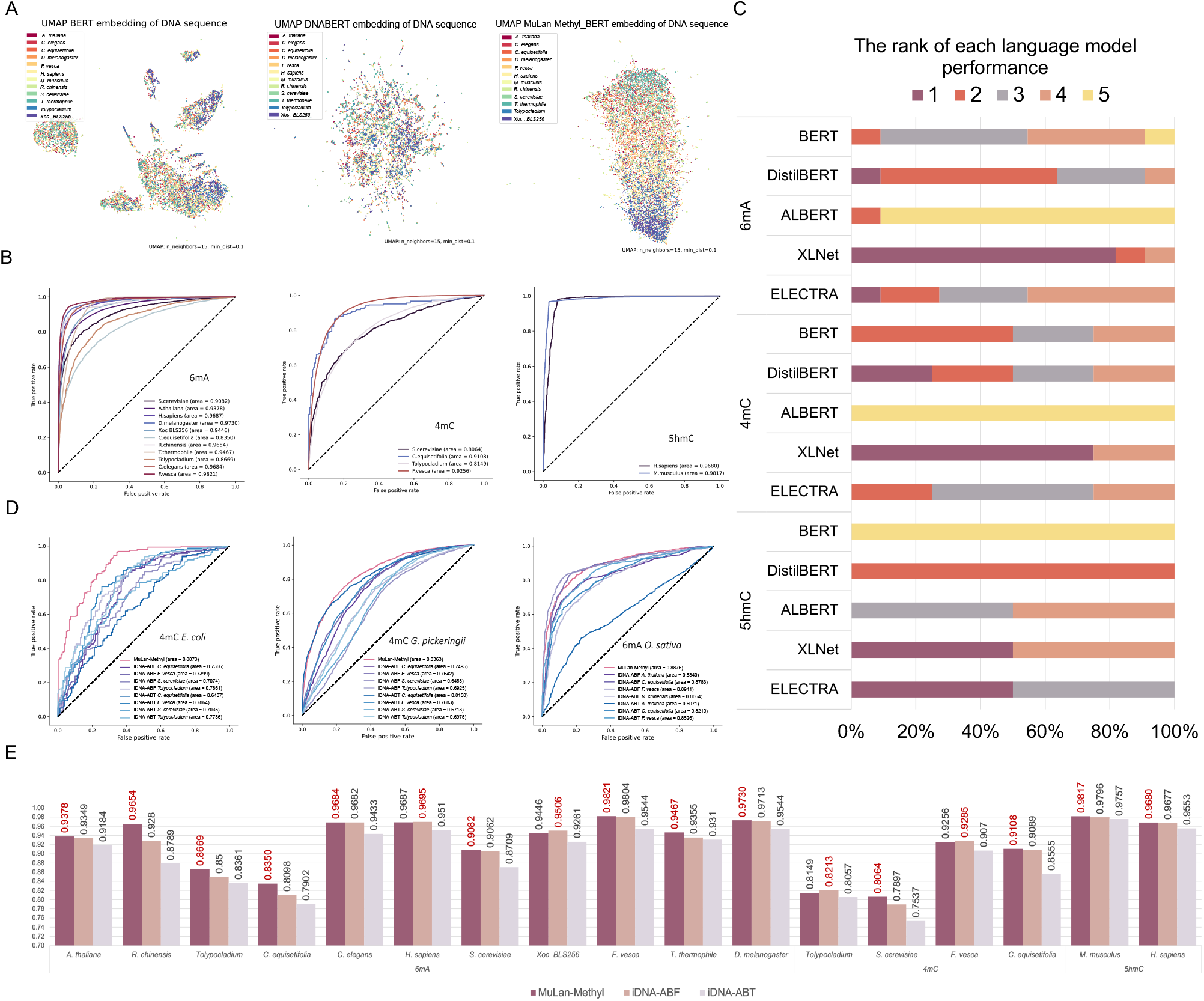
Model analysis and performance comparison of MuLan-Methyl. **(A)** UMAP clustering of sample representations encoded by different pre-trained models; BERT, DNABERT, and MuLan-Methyl BERT (from left to right). Samples are colored by taxonomic lineage. **(B)** For MuLan-Methyl predictions of the three methylation-site types, 6mA, 4mC, and 5hmC, we present ROC curves for each of the 12 taxonomic types in the dataset. The AUC values are shown in brackets. **(C)** For each of the three methylation-site types, and each of the five language models, BERT, DistilBERT, ALBERT, XLNet, and ELECTRA, we show the ranking of models over all taxonomic lineages in terms of AUC scores. Moreover, the frequency with which each fine-tuned model appeared in indicated as the width of the corresponding block. **(D)** Comparion of MuLan-Methyl against two published methods, iDNA-ABF and iDNA-ABT, on an additional dataset that only contains taxonomic lineages that were not used to train the methods. From left to right, we show the ROCs obtained for the prediction of 4mC-sites in *E. coli*, 4mC-sites in *G. pickeringiin* data, and 6mA-sites in *O. sativa* L. data, respectively. **(E)** Comparison of MuLan-Methyl against iDNA-ABF and iDNA-ABT, on the iDMA-MS independent test set. We display the AUC scores for all three methods, for each of the three methylation-site types and each of the 12 taxonomic lineages.

Since the original corpus that BERT is trained on does not explicitly includes DNA fragments, during tokenization, BERT will represent each DNA 6-mer with the special symbol [UNK], or cuts it into small pieces, unaware that it is a biological sequence. Consequently, the DNA sequences are embedded into a sparse space distribution by this encoder, with a poor ability to distinguish different species.

DNABERT is trained on genome sequences and has a better ability to capture DNA sequence features, as reflected in the absence of significant gaps between the distribution of DNA sequence representation obtained by its encoder. In the UMAP visualization, colors representing different taxonomic lineages appear to be randomly distributed.

In comparison, the MuLan-Methyl-BERT encoder is better at identifying DNA fragments and differentiating sequences by taxonomic lineage. Colors representing different taxonomic lineages exhibit a gradient from top to bottom. This suggests that pre-training the language model using a custom corpus that contains both DNA 6-mers and taxonomic lineages, signficantly improves the models ability to capture potential information in this application scenario.

### Comparison with single language sub-models

The MuLan-Methyl framework uses five language models. In this section, we establish that the average prediction probability of this integrated approach is better than using any of the individual sub-models, by comparing model performance using AUC values. In summary, MuLan-Methyl outperforms the sub-models, displaying the highest AUC across different taxonomic lineages and for each methylation-site type.

In more detail, for 6mA-site prediction, MuLan-Methyl is most beneficial when predicting on *Tolypocladium* and *S. cerevisiae*, with an AUC gain of 1.4% over the AUC calculated by ALBERT, which was the best-performing sub-model. The average increase of AUC compared to the taxonomic-lineage-specific best sub-model is 0.7%. For 4mC-site prediction, the average gain of AUC computed from MuLan-Methyl is 0.8%, where the biggest improvement using MuLan-Methyl happened on *S. cerevisiae*, with an AUC increase of 1.1% over XLNet, the best sub-model for this taxonomic lineage. MuLan-Methyl performs as well as taxonomic-lineage-specific sub-models at identifying 5hmC-sites on both of the genomes. Moreover, we assessed the performance of MuLan-Methyl for each methylation-site type and report on this for each taxonomic lineage using multiple metrics, including accuracy, F1-score, recall and precision, and AUC (see Table 1, Table 2, Table 3), as well as their ROC curve (see Figure 2B).

**Table 1.** MuLan-Methyl prediction performance on 6mA-sites.

| Lineage | AUC | Accuracy | F1 | Recall | AUPR |
| --- | --- | --- | --- | --- | --- |
| *T. thermophile* | 0.9467 | 0.8840 | 0.8923 | 0.9611 | 0.9321 |
| *A. thaliana* | 0.9378 | 0.8649 | 0.8615 | 0.8401 | 0.9423 |
| *H. sapiens* | 0.9687 | 0.9077 | 0.9068 | 0.8975 | 0.9721 |
| *Xoc. BLS256* | 0.9446 | 0.8742 | 0.8712 | 0.8511 | 0.9421 |
| *D. melanogaster* | 0.9730 | 0.9276 | 0.9275 | 0.9258 | 0.9761 |
| *C. elegans* | 0.9684 | 0.9131 | 0.9138 | 0.9219 | 0.9674 |
| *C. equisetifolia* | 0.8350 | 0.7590 | 0.7481 | 0.7158 | 0.8492 |
| *S. cerevisiae* | 0.9082 | 0.8325 | 0.8233 | 0.7802 | 0.9198 |
| *Tolypocladium* | 0.8669 | 0.7895 | 0.7824 | 0.7567 | 0.8730 |
| *F. vesca* | 0.9821 | 0.9407 | 0.9403 | 0.9336 | 0.9831 |
| *R. chinensis* | 0.9654 | 0.9164 | 0.9167 | 0.9197 | 0.9691 |

**Table 2.**
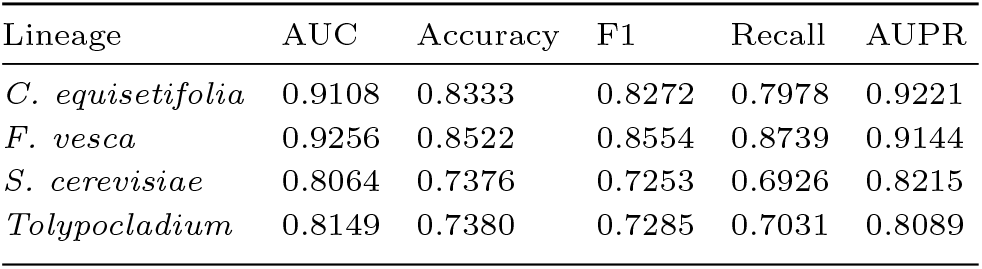
MuLan-Methyl prediction performance on 4mC-sites.

| Lineage | AUC | Accuracy | F1 | Recall | AUPR |
| --- | --- | --- | --- | --- | --- |
| *C. equisetifolia* | 0.9108 | 0.8333 | 0.8272 | 0.7978 | 0.9221 |
| *F. vesca* | 0.9256 | 0.8522 | 0.8554 | 0.8739 | 0.9144 |
| *S. cerevisiae* | 0.8064 | 0.7376 | 0.7253 | 0.6926 | 0.8215 |
| *Tolypocladium* | 0.8149 | 0.7380 | 0.7285 | 0.7031 | 0.8089 |

**Table 3.** MuLan-Methyl prediction performance on 5hmC-sites.

| Lineage | AUC | Accuracy | F1 | Recall | AUPR |
| --- | --- | --- | --- | --- | --- |
| *M. musculus* | 0.9817 | 0.9649 | 0.9651 | 0.9685 | 0.9782 |
| *H. sapiens* | 0.9680 | 0.9484 | 0.9500 | 0.9787 | 0.9485 |

For each of the three methylation-site types, and for each of the five sub-models included in MuLan-Methyl, we evaluated the performance of sub-models on the corresponding independent test set. For each of the 12 taxonomic lineages, we ranked the give sub-models based on their AUC values. Also, we determined the occurrence frequency of each sub-model at each rank. This is shown in Figure 2C and Supplementary Figure S1.

We observed that XLNet usually shows better AUC than the other sub-models for predicting 6mA-sites, ranked first for 9 lineages. In contrast, ALBERT performs very poorly.

XLNet also performed best in 4mC-site predictions, achieving the highest AUC on 3 out of 4 taxonomic lineages. The lowest AUC from 4 taxonomic types concentratedly results from ALBERT. XLNet and ELECTRA performed best on 5hmC-site, BERT performs worst.

### Comparison with existing methods

To demonstrate the advantage of MuLan-Methyl over existing methods, we compared the method against iDNA-ABF and iDNA-ABT, two state-of-the-art methods, that are both able to predict methylation-sites for all three types, across different taxonomic lineages. (Note that all three frameworks were trained on the same training dataset, provided by iDNA-MS.) For this, we used the iDNA-MS independent test set, which is considered a benchmark dataset. We report the AUC scores in Figure 2E, and more comprehensive evaluation metrics are displayed in Supplementary Table S2.

In this study, MuLan-Methyl outperforms the other two methods on 13 out of 17 combinations of methylation types and taxonomic lineages. First, for 6mA-site prediction, MuLan-Methyl improves over the other methods by between 0.02% to 3.74% AUC, whereas for *R. chinensis, C. equisetifolia, Tolypocladium*, and *T. thermophile*, the improvement is by more than 1%. Second, for 4mC-site prediction, our method shows an increase of 1.67% and 0.02% AUC, on *S. cerevisiae* and *C. equisetifolia*, respectively. Finally, for 5hmC-site prediction, our method shows an increase of 0.21% and 0.03% on *M. musculus* and *H. sapiens*, respectively.

The iDNA-ABF method has higher AUC scores in the remaining four cases, namely for 6mA-site prediction on *H. sapiens* and *Xoc. BLS256*, with an improvement of 0.08% and 0.6%, and for 4mC-site prediction on *Tolypocladium* and *F. vesca*, with an improvment of 0.64%, and 0.3%, respectively, over MuLan-Methyl. A cursory comparison suggests that MuLan-Methyl and iDNA-ABF have similar reported run times (albeit using different GPUs), whereas iDNA-ABT runs about ten times faster.

### Explainability of MuLan-Methyl aids motifs discovery

To assess the contribution of each token toward correct methylation-site detection, we use the average attention weight assigned by each token to [CLS] in the fine-tuned sub-model, based on the positive sample from the independent test set.

The importance scores of each position in a DNA sequence has a Gaussian distribution across 17 different combinations of methylation-site types and taxonomic lineages (see Figure 3D-F and Supplementary Figure S3). Positions of higher importance are concentrated around the center of the samples, and the central position always has high significance.

**Fig. 3.**
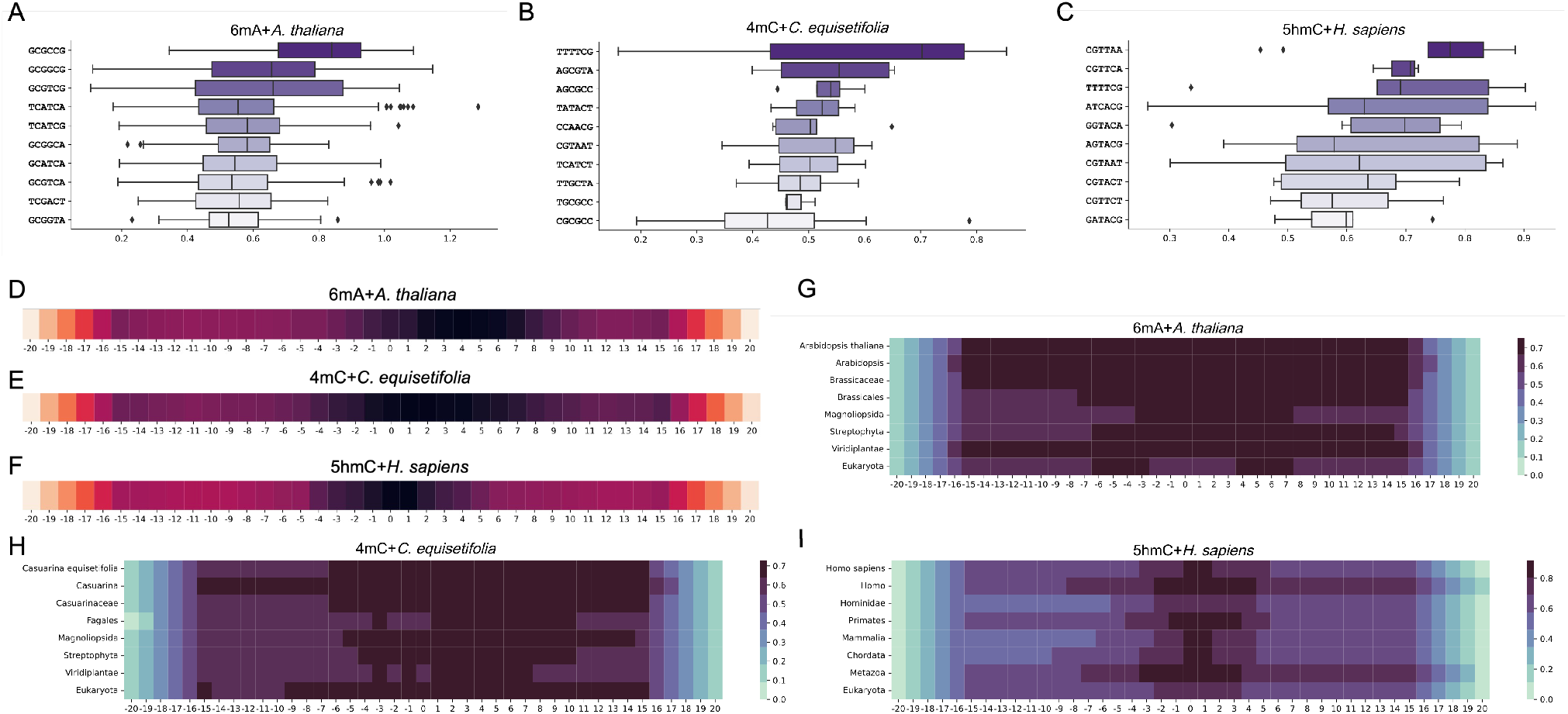
Interpretation of MuLan-Methyl by attention weights resulting from transformer self-attention mechanism. In **(A)**–**(C)**, box-plots show the distribution of attention weights for the ten 6-mer of hightest average importance scores, for the combinations 6mA + *A. thaliana*, 5mC + *C. euisetifolia* and 5hmC + *H. sapiens*, respectively. In **(D)**–**(F)**, we indicate the importance score for each position in the DNA sequences of length 41, obtained by merging 6-mer fragments, for the same three combinations listed above, respectively. In **(G)**–**(I)**, for each taxonomic rank of a lineage, we indicate the attention weight assigned by MuLan-Methyl to each position of the sequence for generating the taxon of the given rank, for the same three combinations listed above, respectively.

This observation supports the rationale used for constructing the iDNA-MS dataset, namely to use, as positive samples, DNA segments of length 41 that are each centered on an experimentally verified methylation site. It also suggests the existence of DNA motifs that are closely associated with DNA methylation.

We also observed, for all 17 combinations, that the importance score starts low and then reaches a local maximum at position *±*15. It then steadily increases from *±*16 to the center of each sample (of length 41). This suggests that 41 is an ideal sample length for methylation detection, neither wasting resources to store unimportant positions, nor missing important sequence.

The 6-mers with high importance may be considered to be DNA-methylation “motifs” (see Figure 3A-C and Supplementary Figure S2). For a fixed taxonomic lineage, the three different methylation-site types each have different motifs. However, for a fixed methylation-type-site, some motifs occur across different taxonomic lineages.

For example, the motif CGAAGT is important for 6mA methylation for several taxonomic lineages, namely *S. cerevisiae, Tolypocladium*, and *Xoc. BLS256*. Note that the former two are eukaryotes, whereas the latter is a bacteria. Moreover, for 5hmC methylation, *H. sapiens* and *M. musculus* share many motifs. Similarly, for 4mC methylation, *C. equisetifolia* and *F. vesca* share many motifs.

### Explainability of MuLan-Methyl reveals relationships between DNA sequence and taxonomic lineage

Integrating DNA sequences with taxonomic lineage as an explicit feature adds information and thus increases detection accuracy. Moreover, during fine-tuned model prediction, the association between DNA sequence and taxonomy can be measured by extracting the attention weights assigned from DNA tokens to the tokens that represent taxonomic lineage (see Figure 3G-I and Supplementary Figure S4).

The impact of DNA sequence on taxonomic lineage varies across the 17 combinations of methylation site types and taxonomic lineages. Generally, sequence locations that determine taxonomic lineage are concentrated around the center of samples, where the discussed DNA methylation-associated motifs are also clustered.

Of the eight taxonomic ranks used to specify taxonomic lineage, the highest (kingdom) and lowest rank (species), in particular, are assigned larger attention weights by a wide range of positions in the sequence.

However, not all combinations follow this rule. For example, the impact of DNA sequence on species is weaker than on genus and family for the combinations 6mA + *D. melanogaster* and 5hmC + *M. musculus*. On combinations 6mA + *R. chinensis*, 6mA + *S. cerevisiae*, 6mA + *C. elegans*, 4mC + *S. cerevisiae*, and 5hmC + *H. sapiens*, we observed that the high scores assigned to the taxonomy lineages are quite sparsely distributed over the different ranks.

These observations demonstrate that the explainability of MuLan-Methyl can shed light on the relationships between DNA sequences and taxonomic lineage.

### Performance on the external dataset

MuLan-Methyl was trained on 17 combinations of DNA methylation-site types and taxonomic lineages. Fine-tuned models aim at performing well on input whose distribution is consistent with the training dataset, however are not guaranteed to perform well on other data.

To explore the performance of MuLan-Methyl on other data, we applied the approach to an external dataset that contains three combinations of methylation types and taxonomic lineages, namely 4mC + *E. coli*, 4mC + *G. pickeringi* and 6mA + *O. sativa* L. Note that these three taxonomic lineages do not appear in the iDNA-MS datasets.

For the sake of comparison, we also calculated predictions using the servers provided by iDNA-ABF and iDNA-ABT. Since both approaches provide independent models for each combination, we run all taxon-wise models for 4mC-site detection, and the appropriate ones for 6mA-site detection.

MuLan-Methyl performed much better than the other two models on the 4mC + *E. coli* combination, achieving an AUC of 0.89, more than 10% better than the others. Our method also performed best on the 4mC + *G. pickeringi* combination, with an advantage of 2.05% over iDNA-ABT (using its *C. equisetifolia* model). On the third combination, 6mA + *O. sativa* L, MuLan-Methyl performed slightly worse (0.65%) than iDNA-ABF (using its *F. vesca* model). See Figure 2D.

### MuLan-Methyl server

We provide an implementation of MuLan-Methyl as a web server (See Fig 4). Like other deep-learning based methylation services, this allows the user to upload DNA samples of length 41, select the closet taxonomic lineage and the type of methylation site of interest. The uploaded sampled will then be classified as methylation sites, or not.

**Fig. 4.**
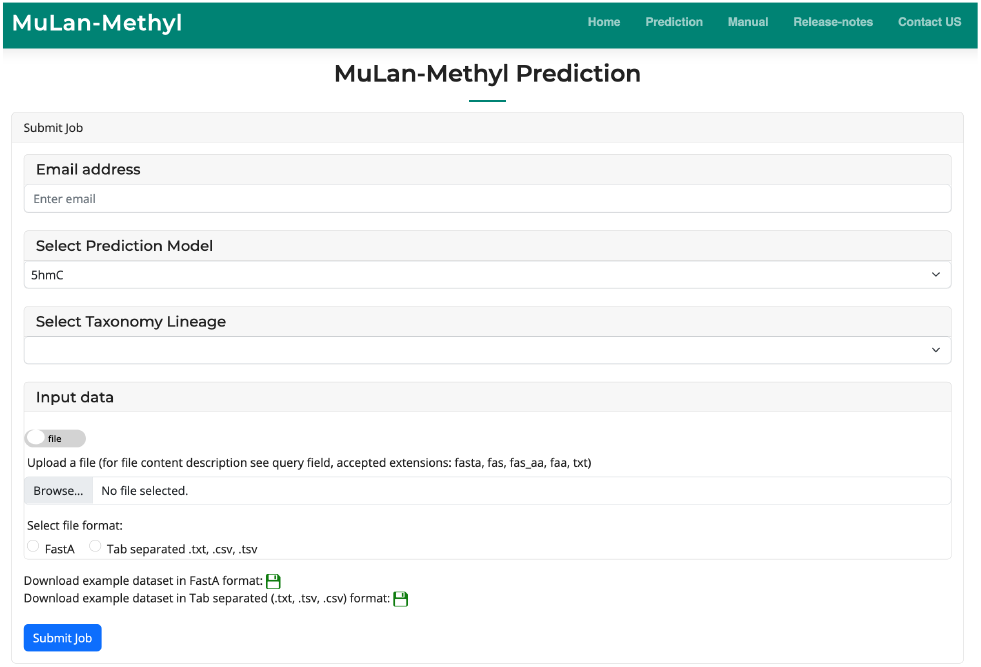
The Mulan-Methyl server hosted at http://ab.cs.uni-tuebingen.de/software/mulan-methyl/ allows upload of DNA sequences and will perform methylation prediction along the whole sequence.

We also allow upload of longer DNA sequences and in this case the server will provide a list of all predicted methylation sites that are predicted in the uploaded sequence.

To implement this extended functionality, we first extract all samples of length 41 that are centered on a nucleotide of the appropriate type (e.g. C when predicting 4mC or 5hmC sites) and then perform Mulan-Methyl prediction on these. The predicted positive samples are then filtered by feature importance analysis to resolve overlapping predictions. In more detail, we only retain samples for which the importance scores are highest at the center of the sample. Output is the list of all predicted methylation positions.

## DISCUSSION and CONCLUSION

Previous studies have focused on adapting BERT to specific biological tasks using the pre-train and fine-tune paradigm, with the aim of applying this popular NLP approach to tasks in genomics, phylogenetics and other areas of computational biology.

However, BERT is not the only transformer-based language model and it is important to choose the best model for a given task. Our proposed framework MuLan-Methyl consists of five transformer-based language models for identifying three types of DNA methylation sites across several taxonomic lineages, including both Eukaryota and Bacteria. With this work, we extend the list of transformed-based language models that have been successfully adapted to tasks involving biological sequences.

Each sub-model in MuLan-Methyl is pre-trained and fine-tuned on the training dataset, and they then collectively predict methylation sites on an independent test dataset. The performance of MuLan-Methyl was evaluated by multiple metrics and in comparison with two existing approaches, and the method showed very good performance.

Our study also indicates that models with enhanced algorithms in the pre-training step, such as XLNET, and models with fewer parameters and less memory consumption, such as DistilBERT, are more appropriate than BERT when storage or computational resources are limited.

In contrast to other biological domain-adaption language models, the custom corpus that we trained MuLan-Methyl on contains multi-modal data, consisting of both DNA sequences from iDNA-MS and taxonomy lineage in text format from the NCBI and GTDB taxonomies. To the best of our knowledge, MuLan-Methyl is the first language-model framework to take taxonomy information into consideration.

This improves model accuracy and feature contribution analysis. The DNA methylation motifs found by MuLan-Methyl greatly benefited from the self-attention mechanism of transformer structure. In addition, the attention weights assigned to taxonomic lineages by DNA sequences help to analyze the relationship between nucleotide sequences and taxonomy lineage.

Previous approaches build a separate classifier for each taxonomic lineage and each methylation-site type, giving rise to 17 different classifiers, for the data used here. In contrast, MuLan-Methyl considers taxonomic lineage as a feature and so only gives rise to three classifiers, one for each type of methylation site.

This study demonstrates that BERT is not the only choice when one wants to adapt a transformer-based language model to a specific domain, one should also consider its variants. It also shows that integrating multiple language models can offset the deficiencies of the individuals models, to some extent, so as to obtain an improved ensemble prediction performance.

In conclusion, we have proposed a framework that integrates five popular NLP approaches to solve an important biological problem. MuLan-Methyl can be used to detect DNA methylation sites reliably for DNA sequences of 41 bp length from known taxonomic lineages, especially when closely related to the lineages involved in training, with slightly better performance than current state-of-the-art methods.

In practical applications, the input will usually be a chromosome or a set of assembled contigs, and the desired output will be a list of putative methylation sites. To address this, we designed a two-step validation strategy for false positive rate controlling and implemented it in the MuLan-Methyl server.

## Data availability

The benchmark dataset used in this study is accessible via the link http://lin-group.cn/server/iDNA-MS/download.html. The processed dataset used for training MuLan-Methyl and the source code are available at https://github.com/husonlab/mulan-methyl. A web server implementing the MuLan-Methyl approach is freely accessible at http://ab.cs.uni-tuebingen.de/software/mulan-methyl/, see also https://bio.tools/MuLan-Methyl, RRID: SCR 023591.

## Supplementary data

Supplementary data are available online at GigaScience.

## Author contributions statement

W.Z. and D.H.H. conceived the project. W.Z. collected and processed the dataset for the project. W.Z. designed and implemented the architecture and algorithms of MuLan-Methyl, and conducted model analysis. A.G. and W.Z. designed and implemented web-server of MuLan-Methyl. W.Z., D.H.H., and A.G. contributed to the manuscript.

## Acknowledgments

We acknowledge support of the BMBF-funded de.NBI Cloud within the German Network for Bioinformatics Infrastructure (de.NBI) (031A532B, 031A533A, 031A533B, 031A534A, 031A535A, 031A537A, 031A537B, 031A537C, 031A537D, 031A538A).

